# SPIN1 promotes tumorigenesis by blocking the uL18-MDM2-p53 pathway

**DOI:** 10.1101/177881

**Authors:** Ziling Fang, Jun-Ming Liao, Bo Cao, Jun Deng, Kevin D Plummer, Peng Liao, Tao Liu, Shelya X. Zeng, Jianping Xiong, Hua Lu

## Abstract

Ribosomal proteins (RPs) play important roles in modulating the MDM2-p53 pathway. However, less is known about the upstream regulators of the RPs. Here we identify SPIN1 (Spindlin 1) as a novel binding partner of human RPL5/uL18 that is important for this pathway. SPIN1 ablation activates p53, suppresses cell growth, reduces clonogenic ability, and induces apoptosis of cancer cells by sequestering uL18 in the nucleolus, preventing it from interacting with MDM2, and thereby alleviating uL18-mediated inhibition of MDM2 ubiquitin ligase activity towards p53. SPIN1 deficiency increases ribosome-free uL18 and uL5 (human RPL11), which are required for SPIN1 depletion-induced p53 activation. Analysis of cancer genomic databases suggests that SPIN1 is highly expressed in several human cancers, and its overexpression is positively correlated with poor prognosis in cancer patients. Altogether, our findings reveal that the oncogenic property of SPIN1 is highly attributed to its negative regulation of uL18, leading to p53 inactivation.

## Introduction

The well-documented tumor suppressor p53, referred as “the guardian of the genome”, is activated upon exposure to a myriad of cellular stresses. While loss of wild type p53 causes fatal damages to cells, it is not surprising that the TP53 gene is mutated in more than 50% human cancers, and the functions of p53 are often impeded through various mechanisms in the remainder ^1^. One predominant negative regulator of p53 is the E3 ubiquitin ligase MDM2, an oncoprotein that conceals the N-terminal transcriptional activation (TA) domain of p53 ^2^ and deactivates this protein by either abrogating its transcriptional activity, or inducing its nuclear export and ubiquitination ^2-5^. A plethora of cellular stress could stabilize p53 by blocking the MDM2-p53 feedback loop ^6^. For example, p19^ARF^ inhibits MDM2-mediated p53 ubiquitination and proteolysis by associating with MDM2 ^7^.

Another pathway is so-called the ribosomal proteins (RPs)-MDM2-p53 pathway ^8,9^. Accumulating evidence has continuingly verified this pathway as an emerging mechanism for boosting p53 activation in response to ribosomal stress or nucleolar stress over the past decade ^10-14^. Ribosomal stress is often triggered by aberrant ribosome biogenesis caused by nutrient deprivation, inhibition of rRNA synthesis or malfunction of ribosomal proteins or nucleolar proteins ^8-11,15,16^. Earlier studies showed that disruption of ribosomal biogenesis induces translocation of a series of ribosomal proteins, including uL18 (human RPL5), uL5 (human RPL11), uL14 (human RPL23), eS7 (human S7) and uS11 (human S14) ^17^, from the nucleolus to the nucleoplasm and bind to MDM2, blocking its ability to ubiquitinate p53 and consequently stabilizing p53 to maintain cellular homeostasis ^12,18-23^. Although there are a few proteins that have been identified to regulate this RPs-MDM2-p53 pathway, such as PICT-1 inhibition of uL5 24,25 and SRSF1 inhibition of uL18 ^26^, it still remains to determine if there are more proteins that can regulate the RPs-MDM2-p53 pathway. In this present study, we identified SPIN1 as another uL18 regulator.

SPIN1, a new member of the SPIN/SSTY family, was originally identified as a highly expressed protein in ovarian cancer ^27^. The oncogenic potential of SPIN1 was later supported by the observation that overexpression of SPIN1 increases transformation and tumor growth ability of NIH3T3 cells ^28^. Signaling pathways responsible for SPIN1 functions include PI3K/Akt, Wnt and RET that are highly relevant to tumorigenesis ^29-31^. In addition, SPIN1 acts as a reader of histone H3K4me3 and stimulates the transcription of ribosomal RNA-encoding genes ^32-34^, suggesting its role in rRNA synthesis.

In screening uL18-associated protein complexes using co-immunoprecipitation followed by mass spectrometry, we identified SPIN1 as one of the potential uL18 binding proteins. We confirmed the specific interaction of SPIN1 with uL18, but not with uL5 or uL14, and also found out that by binding to uL18, SPIN1 prevents the inhibition of MDM2 by uL18 and promotes MDM2-mediated p53 ubiquitination and degradation. Also, SPIN1 knockdown induced ribosomal stress by facilitating the release of ribosome-free uL18 or uL5, accompanying p53 activation. Furthermore, SPIN1 knockdown inhibited cell proliferation and induced apoptosis in a predominant p53-dependent manner in vitro and in vivo, consequently suppressing tumor growth in a xenograft model. Therefore, these results for the first time demonstrate that SPIN1 can regulate the RP-MDM2-p53 pathway by directly interacting with uL18, and suggest SPIN1 as a potential molecule target in this pathway for developing anti-cancer therapy in the future.

## RESULTS

### SPIN1 interacts with uL18

Our and others’ studies previously demonstrated that uL18 can stabilize p53 by binding to MDM2 and inhibiting its E3 ligase activity toward p53 ^19,35^. In order to identify potential upstream regulators that may modulate the uL18-MDM2-p53 circuit, we performed co-immunoprecipitation (co-IP) using HEK293 cells that stably expressed Flag-uL18 with the anti-Flag antibody, and the co-immunoprecipitated proteins were cut out for mass spectrometric (MS) analysis (Fig. 1A). The MS results not only revealed several previously described p53 regulatory proteins, such as MYBBP1A, PRMT5 and SRSF1, as uL18 binding proteins (Table S1), but also identified SPIN1 as a novel uL18-binding protein candidate that was previously shown to play a role in tumorigenesis and rDNA transcription ^30,34^.

**Figure 1.**
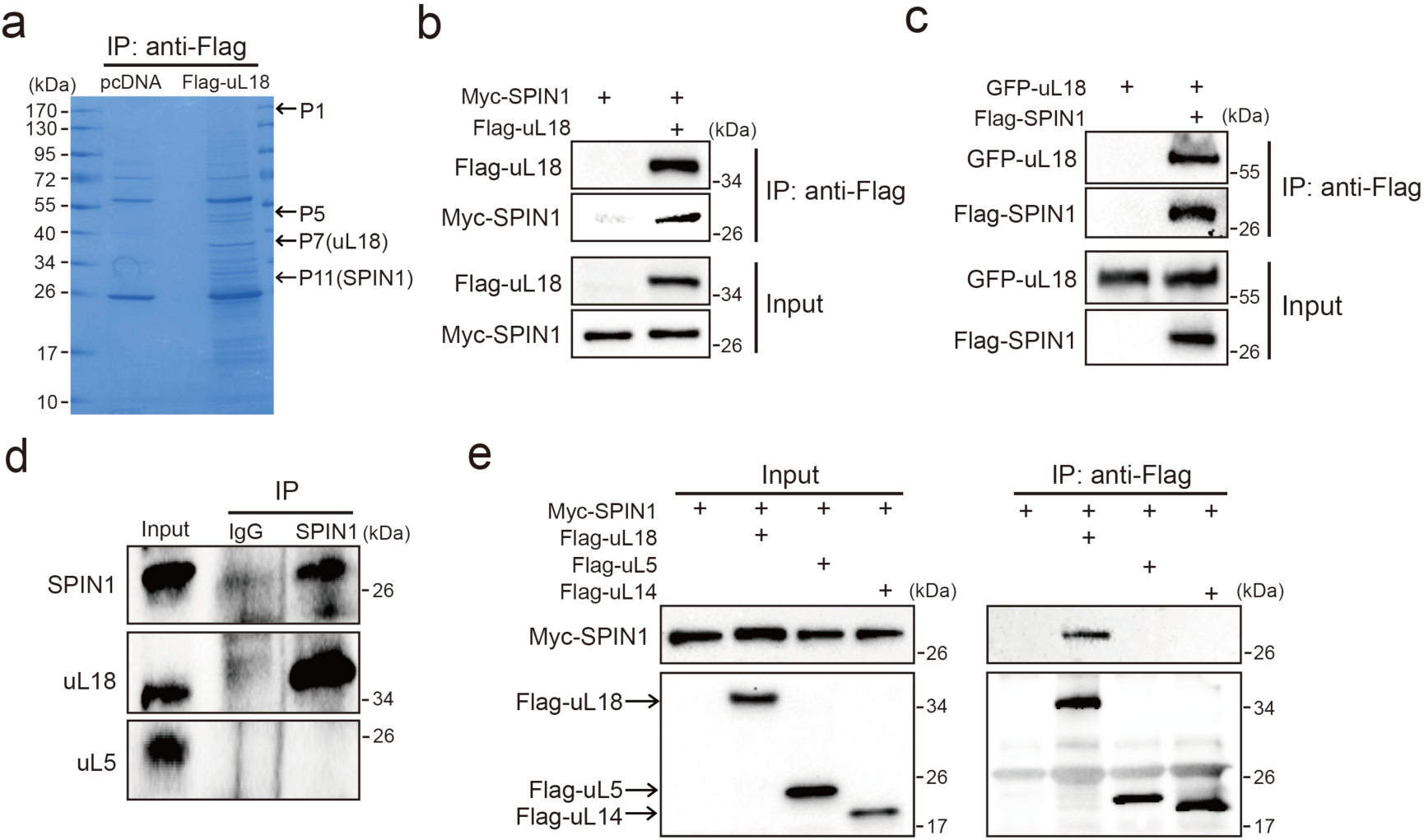
SPIN1 binds to uL18, but not uL5, or uL14. (**a**) Identification of SPIN1 as a candidate of uL18 binding protein by immunopurification and mass spectrometric analysis. Lysates from HEK 293 cells were immunoprecipitated with the anti-Flag antibody. Bound proteins were visualized on a coomassie staining SDS-PAGE gel. Several bands were excised and subjected to mass spectrometry. One of them was identified as SPIN1 (Spindlin 1). The polypeptides identified from these bands are listed in supplementary Table 1. (**b**) and (**c**) SPIN1 interacts with uL18. (**b**) HCT116^p53-/-^ cells were transfected with plasmids encoding Myc-SPIN1 and Flag-uL18, and 48 h later cell lysates were collected for immunoprecipitation (IP) analysis using the anti-Flag antibody. (**c**) HCT116^p53-/-^ cells were transfected with plasmids encoding Flag-SPIN1 and GFP-uL18 for 48 h and harvested for WB analysis with indicated antibodies. (**d**) The interaction between endogenous SPIN1 and uL18. The HEK 293 cell lysates were immunoprecipitated with anti-SPIN1 or control immunoglobulin G (IgG), followed by WB analysis with anti-SPIN1, anti-uL18 and anti-uL5. (**e**) SPIN1 was specifically immunoprecipitated by uL18, but not uL5 or uL14. H1299 cells were co-transfected with Myc-SPIN1 and Flag-uL18, Flag-uL5 or Flag-uL14 as indicated and subjected to IP with the anti-Flag antibody, followed by WB analysis with indicated antibodies.

Next, we confirmed the interaction between SPIN1 and uL18 by performing a series of reciprocal co-IP assays. As expected, ectopic SPIN1 was specifically pulled down by ectopic uL18 and vice versa in HCT116^p53-/-^ cells (Fig. 1B and 1C). Their interaction was also verified in HEK293 cells (Supplementary Fig. S1). Also, we validated the interaction between endogenous SPIN1 and uL18 in HEK293 cells using anti-SPIN1 antibody (Fig. 1D). Interestingly, only uL18, but not uL5, was co-immunoprecipitated with SPIN1. In line with this result, when comparing ectopic Flag-uL18 with Flag-uL5 and Flag-uL14, we found that only uL18, but not the other RPs, could pull down Myc-SPIN1 (Fig. 1E), further bolstering the specific interaction between uL18 and SPIN1. Taken together, these results demonstrate that SPIN1 specifically binds to uL18, but not uL5 or uL14, in cancer cells.

### SPIN1 knockdown inhibits proliferation and induces apoptosis of cancer cells by activating p53

Previous and recent studies showed that SPIN1 is a potential oncogene ^29,30,36^, and uL18 can stabilize p53 by binding to MDM2 ^19^. We therefore wondered if the interaction between SPIN1 and uL18 could confer any role to SPIN1 in regulation of the p53 pathway. First, we determined if depletion of SPIN1 might affect p53-dependent cellular outcomes. Interestingly, we found that knockdown of SPIN1 dramatically elevates p53 protein level in several wild type p53-containing cells, including U2OS, H460 and HCT116^p53+/+^ cells (Fig. 2A), without affecting p53 mRNA expression (Fig. 2B). Consistently, protein and mRNA levels of p53 target genes, such as p21 and PUMA, were also increased in response to SPIN1 knockdown (Fig. 2A and 2B). Moreover, the effects of SPIN1 siRNA on p53 activity were dose-dependent (Supplementary Fig. S2). Conversely, overexpression of SPIN1 in HCT116^p53+/+^ decreased the protein levels of p53 and its targets, such as p21 and PUMA, and the mRNA levels of these target genes, without affecting p53 mRNA level (Fig. 2C and 2D).

**Figure 2.**
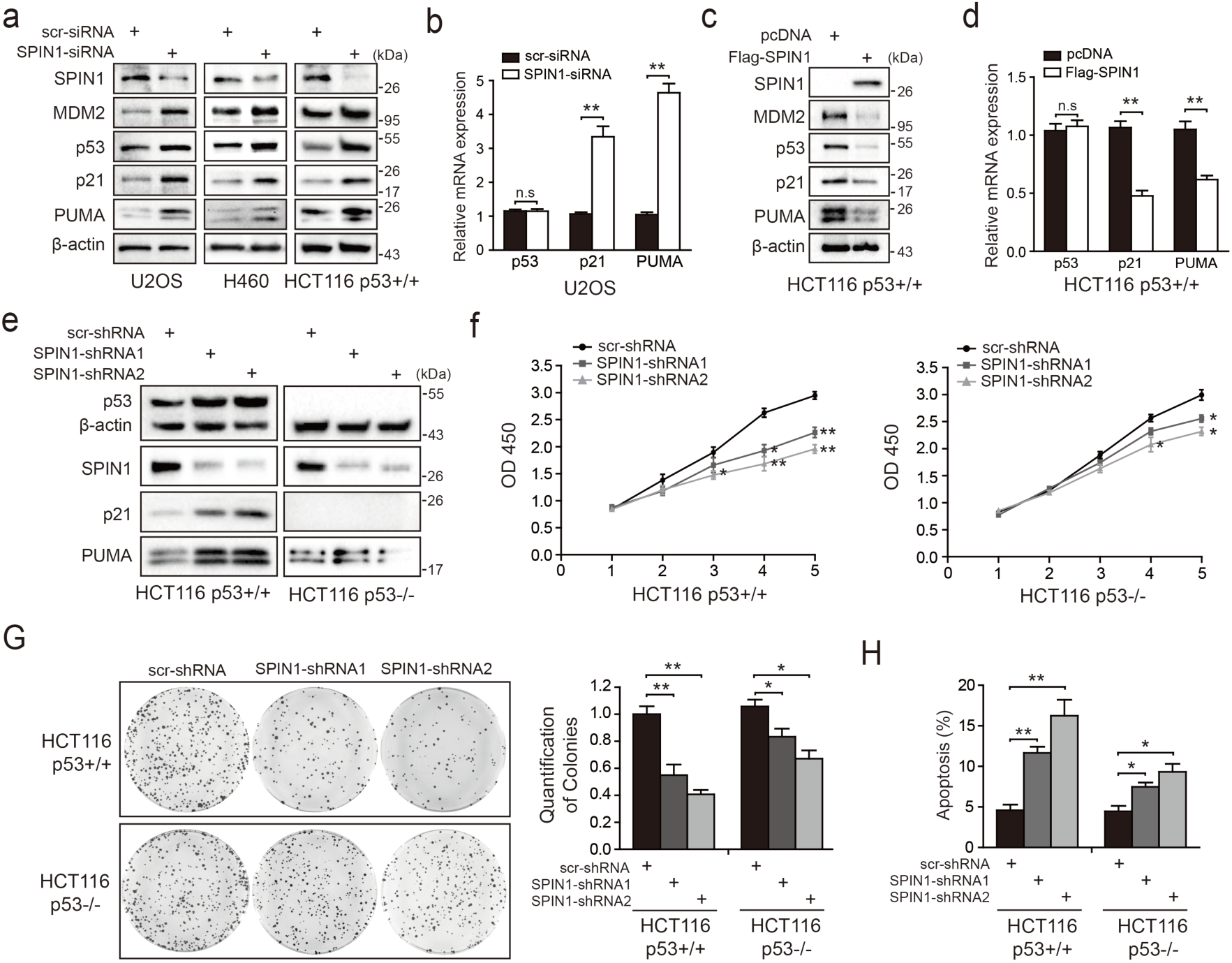
SPIN1 knockdown inhibits cell proliferation and induces apoptosis. (**a**) SPIN1 knockdown induces protein levels of p53 and its target genes. U2OS, H460 and HCT116^p53+/+^ cells were transfected with scramble siRNA (scr-siRNA) or SPIN1 siRNA and harvested 48 h post-transfection for WB analysis with indicated antibodies. (**b**) SPIN1 knockdown induces mRNA levels of p53 target genes without effect on p53 mRNA level. U2OS cells were transfected with scramble siRNA (scr-siRNA) or SPIN1 siRNA, and harvested 72 h post-transfection for RT-qPCR (mean ± SEM, n=2). (**c**) SPIN1 overexpression reduces protein levels of p53 and its target genes. HCT116^p53+/+^ cells were transfected with pcDNA or Flag-SPIN1 and harvested 48 h post-transfection for WB analysis with indicated antibodies. (**d**) SPIN1 overexpression reduces mRNA levels of p53 target genes without effect on p53 mRNA levels. HCT116^p53+/+^ cells were transfected with pcDNA or Flag-SPIN1 and harvested 72 h post-transfection for RT-qPCR (mean ± SEM, n=2). (**e**) Knockdown of SPIN1 causes p53-dependent induction of p21 and PUMA. The protein levels of p53 and its targets in HCT116^p53+/+^ cells and HCT116^p53-/-^ cells that stably express scramble shRNA (scr-shRNA) or SPIN1 shRNAs were detected by WB analysis with indicated antibodies. (**f**) SPIN1 knockdown suppresses cell survival. HCT116^p53+/+^ and HCT116^p53-/-^ cells that stably expressed scramble or SPIN1 shRNAs were seeded in 96-well plate and cell viability was evaluated every 24 h by CCK-8 assays (mean ± SEM, n=2). (**g**) Knockdown of SPIN1 inhibits clonogenic ability of colorectal cancer cells, more significantly when the cells harbor wild type p53. HCT116^p53+/+^ cells and HCT116^p53-/-^ that stably expressed scramble or SPIN1 shRNAs were seeded on 60 mm plates. Puromycin selection was performed for 14 days. Colonies were fixed with methanol, and visualized by staining with crystal violet (mean ± SEM, n=3). (**h**) The effect of SPIN1 knockdown on apoptosis of HCT116^p53+/+^ cells and HCT116^p53-/-^ that stably expressed scramble or SPIN1 shRNAs (mean ± SEM, n=3). \**p*<0.05, \*\**p*<0.01 by two-tailed *t*-test (c, d, g, h).

We next generated both HCT116^p53+/+^ and HCT116^p53-/-^ cell lines that express scramble shRNA or SPIN1 shRNA to evaluate biological outcomes of SPIN1 knockdown. As illustrated in Fig. 2E, the expression of p53 and some of its target genes were markedly induced when SPIN1 was knocked down by its specific shRNA in HCT116^p53+/+^cells, but not in HCT116^p53-/-^cells. Using these cell lines for cell viability assays, we observed that SPIN1 ablation more dramatically represses the cell viability of HCT116^p53+/+^ than that of HCT116^p53-/-^ cells (Fig. 2F). In line with this observation, SPIN1 depletion also led to more predominant reduction of HCT116^p53+/+^ colonies than that of HCT116^p53-/-^ colonies, though both of the reductions were statistically significant (Fig. 2G). Furthermore, percentage of cells undergoing apoptosis caused by SPIN1 shRNAs was much higher in HCT116^p53+/+^ cells than in HCT116^p53-/-^ cells, as measured by sub-G1 population (Fig. 2H). Notably, these effects were proportional to the efficiency of SPIN1 knockdown by two different shRNAs, indicating that the observed effects are highly related to SPIN1 levels. Collectively, these data suggest that SPIN1 plays an oncogenic role by predominantly inactivating the p53 pathway, although SPIN1 may also possess a p53-independent role in cancer cell growth and survival.

### SPIN1 promotes p53 degradation by enhancing MDM2-mediated ubiquitination

Since SPIN1 knockdown affected only the protein, but not the mRNA, levels of p53 (Fig. 2A-2D), we next sought to determine the underlying mechanism. We first performed a cycloheximide-chase experiment using HCT116^p53+/+^ cells. As shown in Figures 3A and 3B, knockdown of SPIN1 markedly prolonged p53’s half-life from ∼30 mins to ∼60 mins, as compared to scramble siRNA. Inversely, ectopic SPIN1 greatly shortened p53’s half-life, from ∼40 mins to ∼20 mins (Fig. 3C and 3D). To further evaluate the effect of SPIN1 on MDM2-mediated p53 ubiquitination, which is the main mechanism responsible for p53 turnover ^12,19,22,37^, we then performed an in vivo ubiquitination assay by transfecting HCT116^p53-/-^ cells with plasmids indicated in Figure 3E. The results clearly showed that ectopic SPIN1 enhances MDM2-mediated p53 ubiquitination in a dose-dependent manner. Consistently, co-transfection of SPIN1 with MDM2 led to a stronger reduction of p53 protein levels, which was abrogated by proteasome inhibitor MG132 (Fig. 3F). Interestingly, the induction of p53 degradation by SPIN1 was MDM2-dependent, as overexpression of SPIN1 failed to repress ectopic p53 protein expression in p53 and MDM2 double knockout MEF cells (Fig. 3G). Together, these results demonstrate that SPIN1 reduces p53 stability by enhancing MDM2-mediated ubiquitination and degradation.

**Figure 3.**
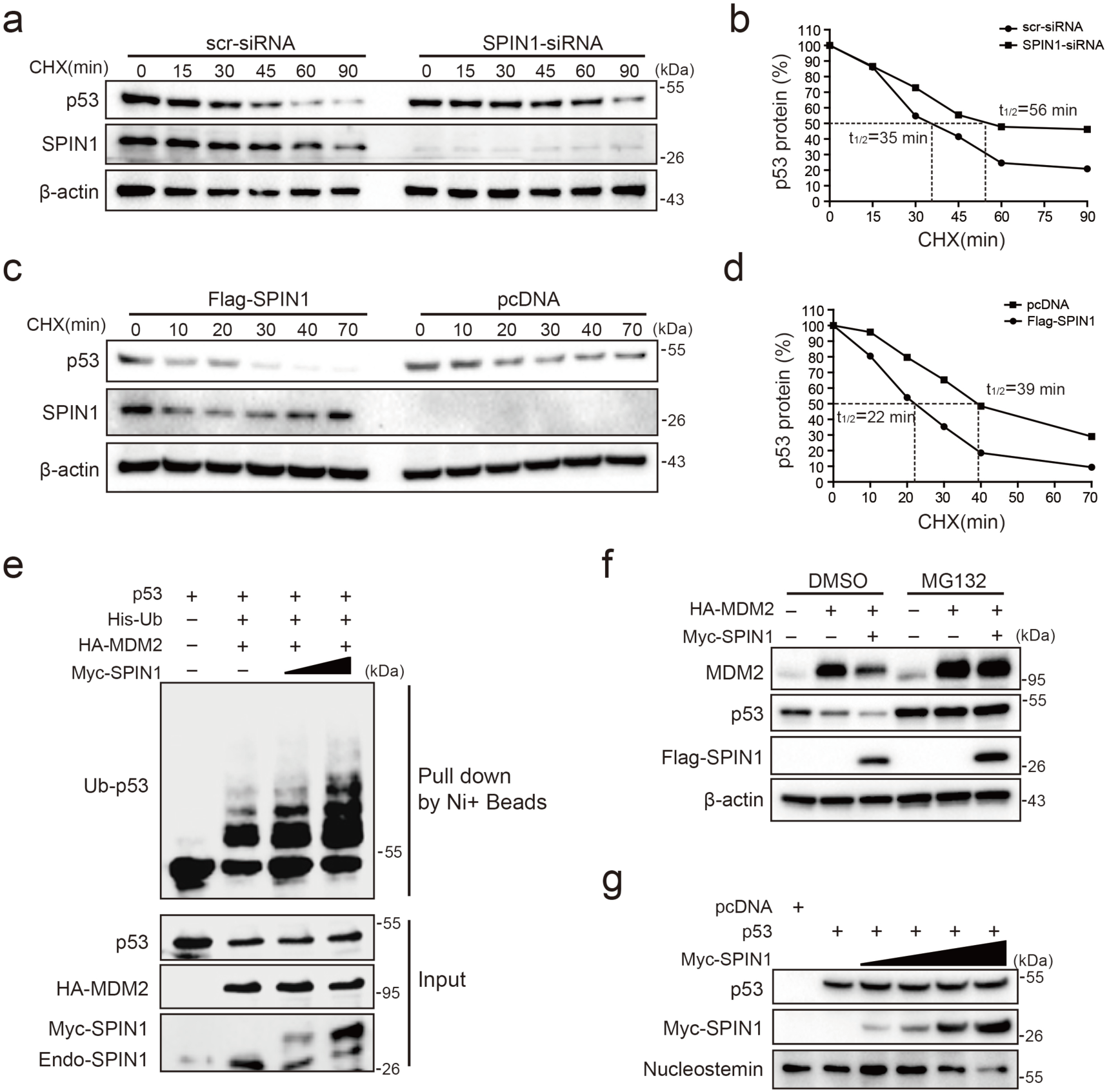
SPIN1 reduces p53 stability by enhancing MDM2-mediated ubiquitination. (**a**) and (**b**) P53-half-life is increased by SPIN1 knockdown. (**a**) HCT116^p53+/+^ cells transfected with scramble or SPIN1 siRNA for 48 h, treated with 100 μg/ml of cycloheximide (CHX), and harvested at different time points as indicated. The p53 protein was detected by WB analysis, quantified by densitometry and plotted against time to determine p53-half-lives (**b**). (**c**) and (**d**) SPIN1 overexpression shortens the half-life of p53. HCT116^p53+/+^ cells transfected with pcDNA or Flag-SPIN1 for 48 h were treated with 100 μg/ml of cycloheximide and harvested at indicated time points for WB analysis with indicated antibodies (**c**). The intensity of each band was quantified, and normalized with β-actin and plotted (**d**). (**e**) SPIN1 promotes MDM2-induced p53 ubiquitination. HCT116^p53-/-^ cells were transfected with combinations of plasmids encoding HA-MDM2, p53, His-Ub or Myc-SPIN1, and treated with MG132 for 6h before being harvested for in vivo ubiquitination assay. Bound and input proteins were detected by WB analysis with indicated antibodies. (**f**) SPIN1 enhances MDM2-mediated p53 proteasomal degradation. HCT116^p53+/+^ cells were transfected with plasmids encoding HA-MDM2 and Flag-SPIN1, and treated with MG132 for 6h before harvested, followed by WB analysis with antibodies as indicated. (**g**) Ectopic SPIN1 does not change p53 protein level without MDM2. MEF^p53-/-; Mdm2-/-^ cells were transfected with combinations of plasmids encoding Myc-SPIN1, HA-MDM2 or p53 followed by WB analysis using antibodies as indicated.

### SPIN1 prevents uL18 from MDM2 binding by sequestering it in the nucleolus

Besides its role as a component of ribosome, uL18 has some well-established extra-ribosomal functions, acting as a bridge in connecting p53 activation to cellular stress response machinery ^8,9^. Upon ribosomal stress, uL18 can translocate from the nucleolus to the nucleoplasm of a cell, where it binds to MDM2 ^19,38^, leading to stabilization of p53 and consequently p53-dependent cell growth arrest, apoptosis or senescence. We then investigated if SPIN1 might regulate this function of uL18, since SPIN1 could bind to uL18 (Figure 1), knockdown of SPIN1 led to p53 activation (Fig. 2), and SPIN1 inhibited MDM2-mediated p53 ubiquitination (Fig. 3). First, as expected ^19^, overexpression of uL18 induced the protein levels of p53 and its targets, such as p21 and MDM2, in wild type p53-containing U2OS cells (Fig. 4A). This induction of the p53 pathway by uL18 was markedly reduced by co-transfected SPIN1 (Fig. 4A). Since the effect of uL18 on p53 is through uL18’s interaction with MDM2 and consequent inhibition of its E3 ligase activity towards p53^19^, we tested if SPIN1 may affect uL18-MDM2 interaction. Interestingly, our co-immunoprecipitation result showed that ectopic Myc-SPIN1 dramatically reduces the amount of Flag-uL18 co-immunoprecipitated with HA-MDM2 in a dose-dependent manner, although Myc-SPIN1 itself did not co-immunoprecipitate with HA-MDM2 (Fig. 4B and S3A). This effect was specific to the uL18-MDM2 interaction, as Myc-SPIN1 overexpression did not alter the interactions between uL5 and MDM2 (Fig. 4C). Our immunofluorescence result (Fig. 4D and S3B) showed that SPIN1 and uL18 are clearly co-localized in the nucleolus, suggesting that SPIN1 might sequester uL18 in the nucleolus and thus prevent it from binding and inactivating MDM2 in the nucleoplasm. Taken together, these results demonstrate that SPIN1 is a regulator of the uL18-MDM2-p53 pathway, acting by preventing uL18 from interaction with MDM2.

**Figure 4.**
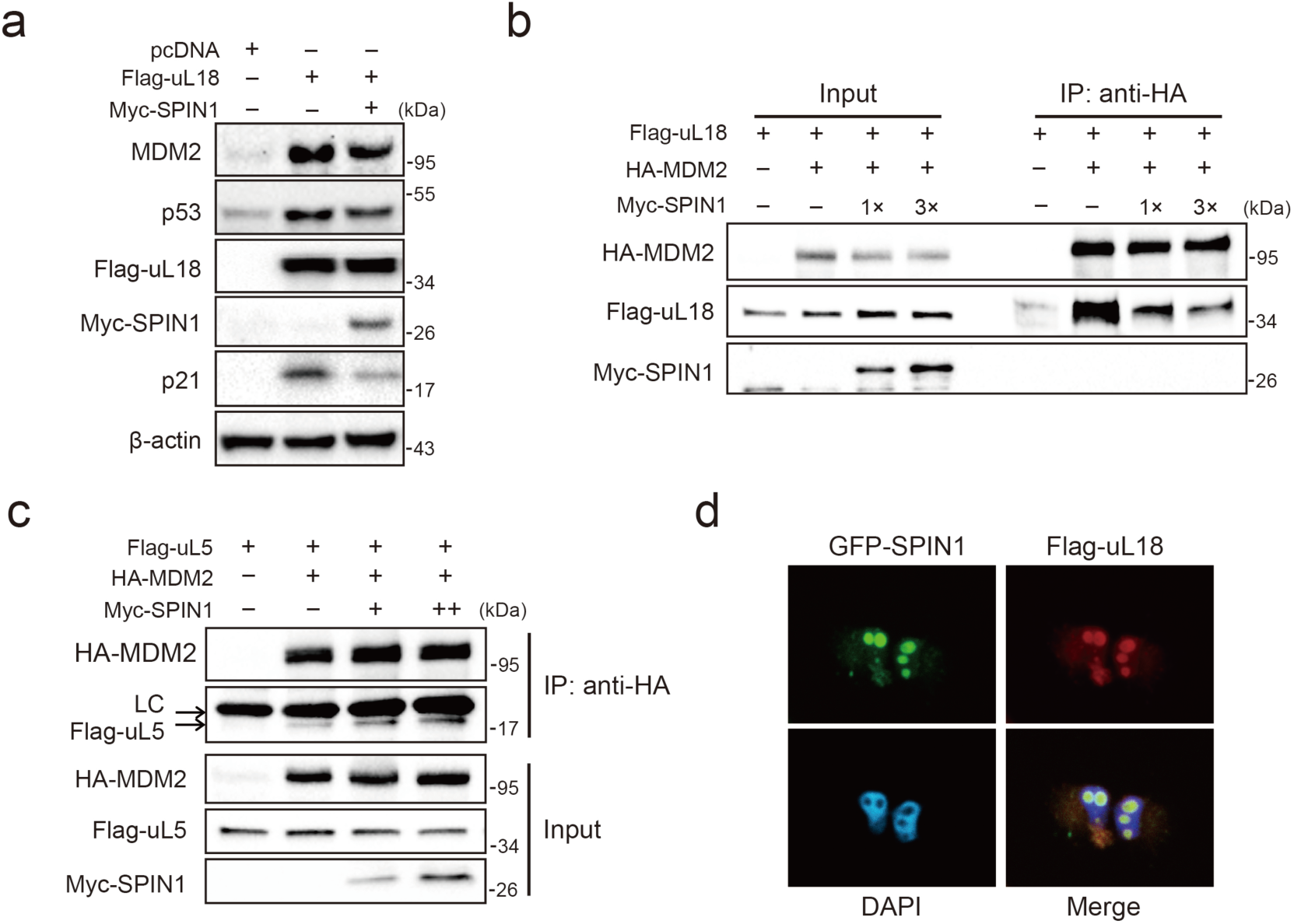
SPIN1 blocks uL18-MDM2 interaction by sequestering uL18 in the nucleolus. (**a**) SPIN1 overexpression attenuates p53 activation induced by ectopic uL18. U2OS cells were co-transfected with plasmids encoding Flag-uL18 or Myc-SPIN1 for 36 h and harvested for WB analysis with indicated antibodies. (**b**) Overexpression of SPIN1 disrupts the uL18-MDM2 binding. Lysates were prepared from HCT116^p53-/-^ cells co-transfected with HA-MDM2, Flag-uL18, Myc-SPIN1 or the corresponding empty vectors for 48 h and analyzed by immunoprecipitated with the anti-HA antibody. Immunoprecipitates and 5% of inputs were immunoblotted with the indicated antibodies. (**c**) Overexpression of SPIN1 fails to disrupt the uL5-MDM2 interaction. Lysates were prepared from HCT116^p53-/-^ cells co-transfected with HA-MDM2, Flag-uL5 and Myc-SPIN1 for 48 h and analyzed by immunoprecipitated with the anti-HA antibody. Immunoprecipitates and 5% of inputs were immunoblotted with the indicated antibodies. (LC: light chain). (**d**) SPIN1 and uL18 co-localize in the nucleolus. H1299 cells were transfected with GFP-SPIN1 and Flag-uL18 for 36 h and then immunostained with the anti-Flag antibody (red), and counterstained with DAPI.

### SPIN1 depletion also causes ribosomal stress, activating p53

Previous studies showed that SPIN1 could recognize H3K4 methylation and stimulate rRNA gene expression, unveiling its role in rRNA synthesis ^32,34^. Disruption of rRNA synthesis leads to disassembly of ribosomal precursors and release of ribosome-free ribosomal proteins from the nucleolus ^16,19,22^. Based on these lines of information, we speculated that dysregulation of SPIN1 itself might also impact ribosome biogenesis, resulting in accumulating ribosome-free ribosomal proteins to activate p53. To test this speculation, we first carried out a sucrose gradient fractionation assay using scramble-and SPIN1-shRNA transfected HCT116^p53+/+^ cells. The collected fractions were subjected to Western blot (WB) analysis. As anticipated, the levels of uL18 and uL5 in the soluble and ribosome-unbound fractions were markedly increased in SPIN1-depletion cells, accompanying with elevated p53 and MDM2 protein levels (Fig. 5A). Interestingly, the binding between endogenous uL18/uL5 and MDM2 increased upon SPIN1 knockdown, resembling ribosomal stress (Fig. 5B). Indeed, as expected, knockdown of SPIN1 reduced the expression of pre-rRNA and rRNA (Supplementary Fig. S4). Moreover, as clearly illustrated in Figure 5C, overexpression of SPIN1 compromised p53 activation induced by actinomycin D or 5-Fu treatment, which was reported to trigger ribosomal stress that in turn triggers the formation of RPs-MDM2 complex ^10,19,23,39^. To further confirm the role of these free forms of ribosomal proteins in SPIN1 ablation-induced p53 activation, we knocked down uL18 or uL5 using siRNA with or without SPIN1 depletion in U2OS cells. Strikingly, the reduction of either uL18 or uL5 abrogated SPIN1 knockdown-induced p53 levels, as well as its target p21, as compared to scramble siRNA-transfected cells (Fig. 5D and 5E). Collectively, these data indicate that knockdown of SPIN1 could also lead to ribosomal stress, releasing ribosome-free uL18 and uL5, which are required for p53 activation induced by SPIN1 depletion.

**Figure 5.**
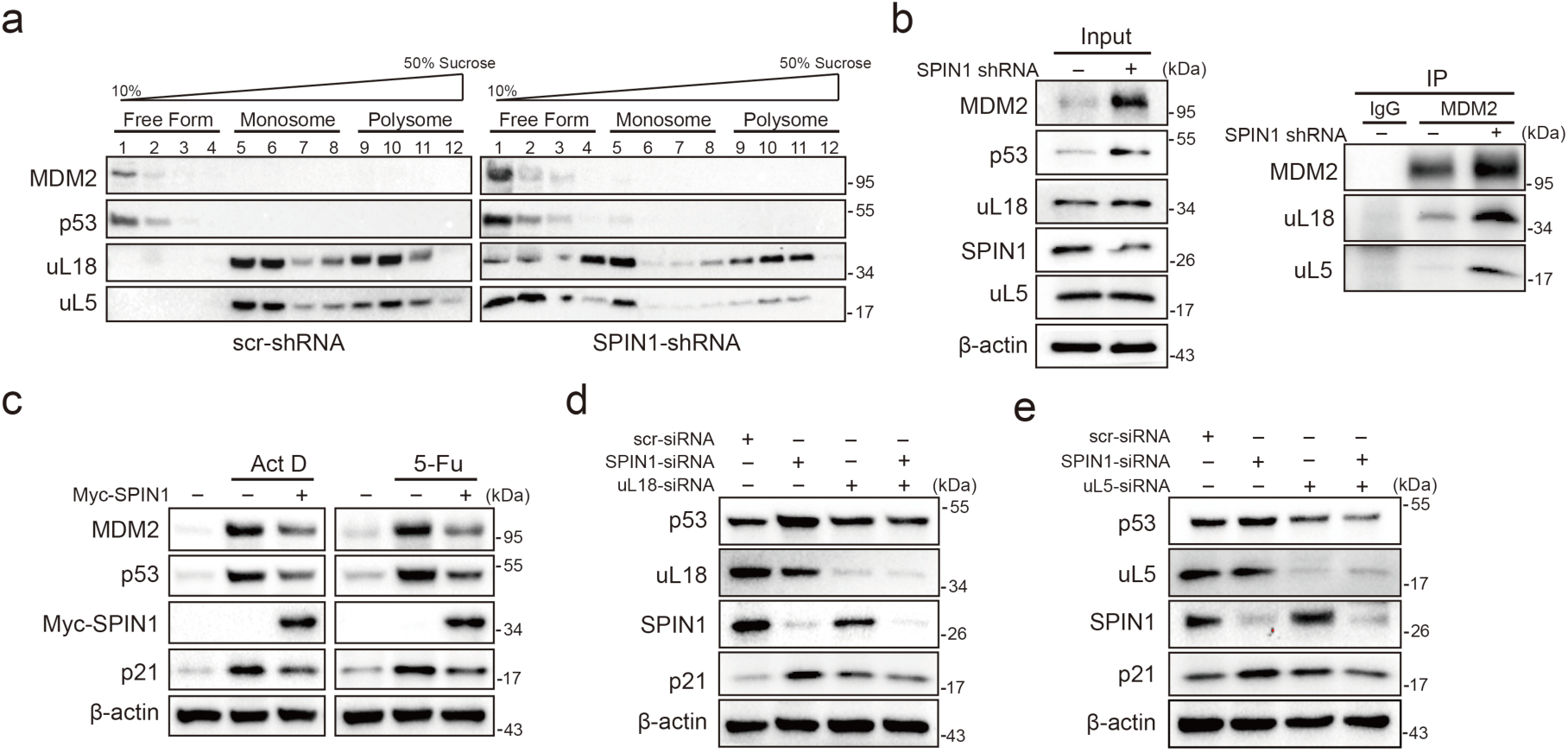
SPIN1 depletion increases ribosome-free uL18 and uL5. (**a**) Knockdown of SPIN1 releases free forms of uL18 and uL5. HCT116^p53+/+^ were transfected with scramble or SPIN1 shRNA for 36 h and subjected to sucrose gradient fractionation assay followed by WB analysis with indicated antibodies. (**b**) SPIN1 knockdown increases the endogenous uL18/uL5-MDM2 interaction. Cell lysates of HCT116^p53+/+^ cells transfected with scramble or SPIN1 shRNA were immunoprecipitated with MDM2 or control IgG, and analyzed by WB analysis with indicated antibodies. (**c**) SPIN1 overexpression counteracts p53 activation induced by ActD or 5-Fu. U2OS cells were transfected with pcDNA or Flag-SPIN1 for 48 h, and treated with ActD or 5-Fu for 12 h before harvested for WB analysis with indicated antibodies. (**d**) and (**e**) Knockdown of uL18 or uL5 compromises the induction of p53 by SPIN1 depletion. U2OS cells were transfected with scramble siRNA, SPIN1 siRNA, uL18 siRNA (**d**) or uL5 siRNA (**e**) as indicated for 48 h. Cell lysates were subjected to WB analysis with indicated antibodies.

### Mapping of uL18- and SPIN1-binding domains

To understand the physical interaction between SPIN1 and uL18 in more details, we generated recombinant GST-SPIN1 and His-uL18 proteins covering full-length or different domains. Our GST-pull down assay showed direct interactions between these two full-length proteins (Fig. 6A and 6C). Additionally, the second Tudor domain of SPIN1 was required for uL18 binding, as His-uL18 was specifically pulled down by the fragments containing the second Tudor domain (amino acid 121-193) (Fig. 6A and 6B). Moreover, both the N- and C-termini of uL18 were found to bind to SPIN1 (Fig. 6C and 6D), and these two fragments were required for uL18-MDM2 binding as well (Supplementary Fig. S5). Based on these data and the aforementioned results, we proposed a model for the role of SPIN1 in regulation of p53 (Fig. 6E). Under the condition of low SPIN1 level, nucleolar uL18 escapes from the nucleolus into the nucleoplasm, and works together with uL5 to bind MDM2 and to inhibit its E3 ubiquitin ligase activity towards p53, consequently leading to p53 activation and p53-dependent cell growth arrest and apoptosis, suppressing cancer cell survival (Fig. 6E, left panel). But when SPIN1 levels are high or abnormally upregulated in cancer cells, SPIN1 retains uL18 in the nucleolus, thereby preventing uL18 from suppression of MDM2 activity and resulting in p53 degradation, favoring tumor cell growth (Fig. 6E, right panel). This conjecture is further supported by the following xenograft experiment.

**Figure 6.**
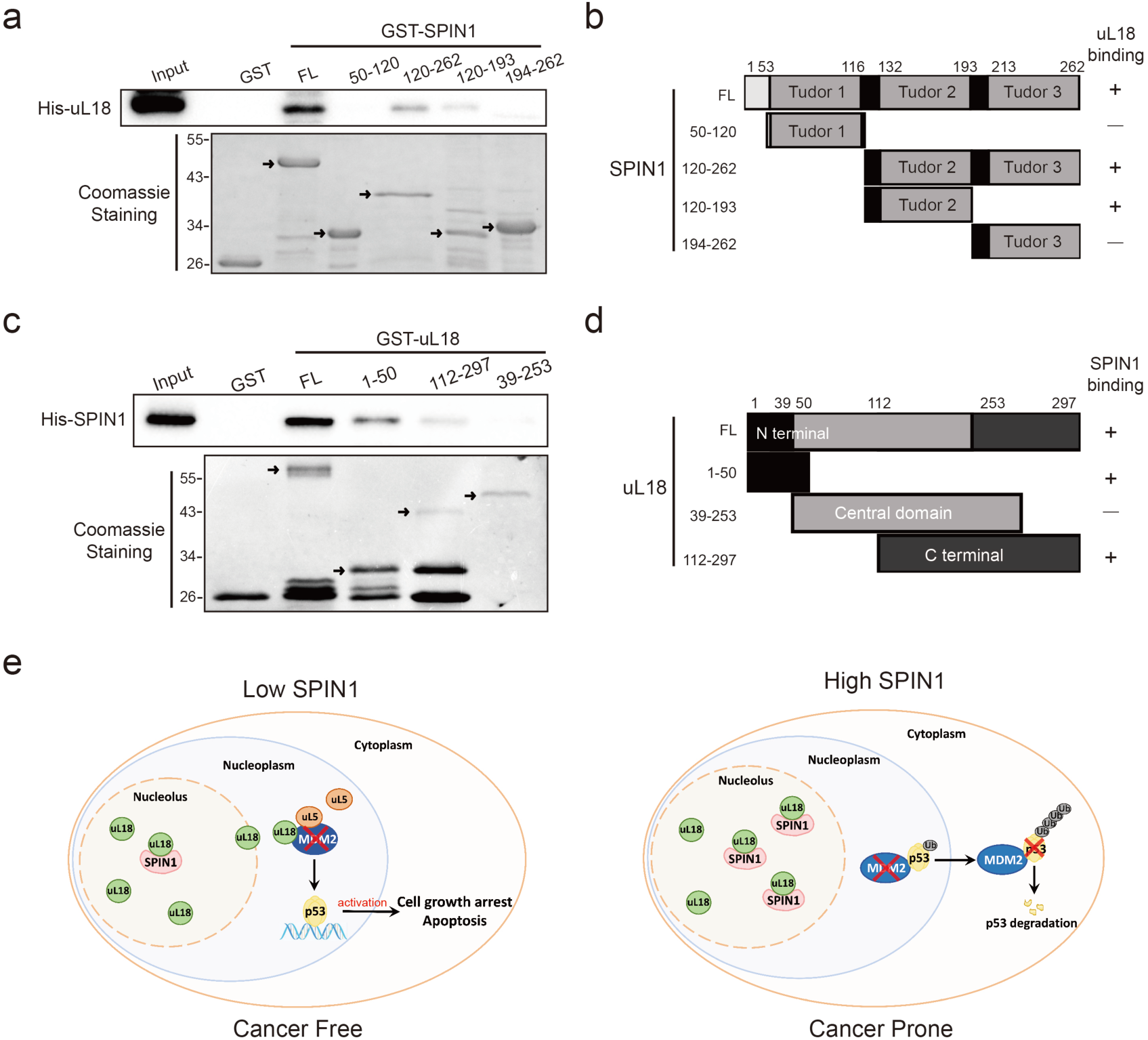
Mapping of uL18- and SPIN1-binding domains. (**a**) uL18 interacts with the second Tudor like domain of SPIN1. Purified GST-tagged SPIN1 fragments, including aa 1-262(FL), aa 50-120, aa 121-262, aa 121-193, aa 194-262 and GST protein alone were incubated with purified His-uL18 protein for 3 hour at 4°C. Bound proteins were detected by WB analysis using anti-uL18 or coomassie staining. (**b**) A schematic diagram of uL18 binding regions on SPIN1 based on the result from (**a**). (**c**) SPIN1 interacts with both the N- and C-termini of uL18. Purified GST-tagged uL18 fragments, including aa1-297(FL), aa 1-50, aa 112-297, aa 39-253 or GST protein alone were rotated with purified His-SPIN1 protein for 1 hour at 4°C. Bound proteins were detected by WB analysis using anti-SPIN1 or coomassie staining. (**d**) A schematic diagram of SPIN1 binding regions on uL18 derived from the result from (**c**). (**e**) A work model for SPIN1 binding to uL18 controls the MDM2-p53 pathway (see text in the Discussion for details).

### SPIN1 depletion impedes xenograft tumor growth

To translate the above-described cellular functions of SPIN1 into more biological significance, we established a xenograft tumor model by inoculating the aforementioned HCT116 (both p53+/+ and p53-/-) cell lines that expressed scramble shRNA or SPIN1 shRNA into NOD/SCID mice, and monitored tumor size for 18 days. As illustrated in Fig. 7A and 7B, SPIN1 knockdown more markedly slowed down the growth of xenograft tumors derived from HCT116^p53+/+^ cells than that from HCT116^p53-/-^ cells. Notably, SPIN1 depletion also reduced the growth of tumors derived from HCT116^p53-/-^ cells, suggesting that SPIN1 might possess a p53-independent function required for cancer cell growth. In line with the tumor growth curve, the reduction of tumor mass and weight by SPIN1 knockdown was more profound in HCT116^p53+/+^ groups (∼60% reduction in weight) than that in HCT116^p53-/-^ groups (∼30% reduction in weight) (Fig. 7C and 7D). To confirm our cell-based findings, we performed qRT-PCR and WB analysis using the xenograft tumors. As expected, the mRNA levels of p21 and PUMA were significantly upregulated upon SPIN1 knockdown in HCT116^p53+/+^, but not in HCT116 ^p53-/-^ tumors (Fig. 7E and 7F). Consistently, the protein levels of p53 and its target PUMA were elevated in HCT116^p53+/+^ groups, but in HCT116^p53-/-^ groups (Fig. 7G and Supplementary Fig. S6). Taken together, these results demonstrate that SPIN1 depletion retards tumor growth by mainly activating p53, though SPIN1 might also possess p53-independent functions in regulation of cell growth and survival.

**Figure 7.**
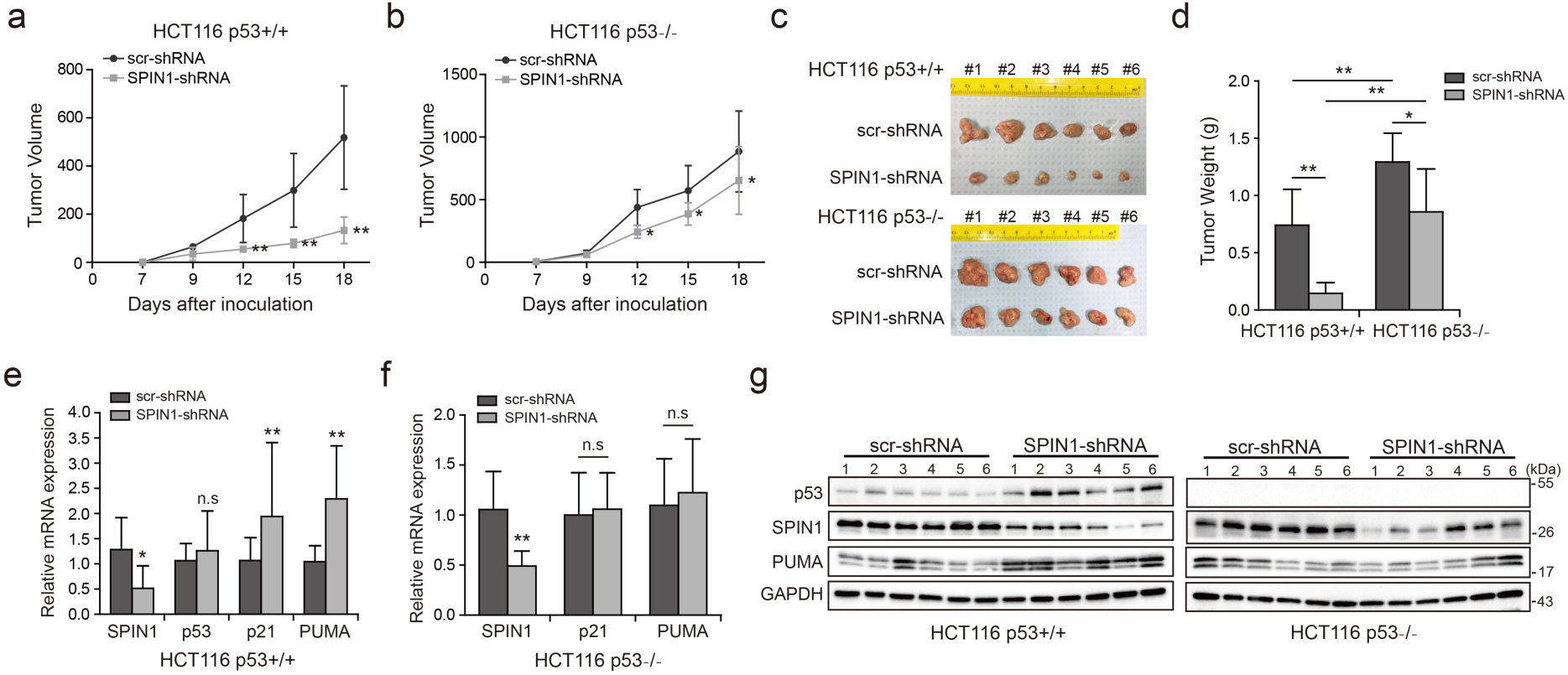
SPIN1 knockdown retards tumor growth more dramatically by inducing p53 activity. (**a**) and (**b**) Growth curves of xenograft tumors derived from HCT116^p53+/+^ cells and HCT116^p53-/-^ cells that expressed scramble or SPIN1 shRNA. Data are represented as mean ± SEM, n=6. (**c**) The images of xenograft tumors that were harvested at the end of experiment. (**d**) Quantification of the average weights of collected tumors from the above experiments. (**e**) and (**f**) The mRNA levels of SPIN1, p53 and p53 target genes were detected in 6 tumors by RT-qPCR (mean ± SEM, n=6). (**g**) The protein levels of SPIN1, p53 and p53 targets were detected in 6 tumors samples by WB analysis with indicated antibodies. \**p*<0.05, \*\**p*<0.01 by two-tailed *t*-test (d, e, f, g).

The data presented above suggest that SPIN1 is required for tumorigenesis. Therefore, we further searched some available genomic and gene expression database for SPIN1 expression in cancers. Interestingly, our analysis of TCGA genome database ^40,41^ indicated that the SPIN1 gene is markedly amplified in a panel of cancers, including prostate, sarcoma, lung, stomach, breast, head and neck, pancreas and colorectal cancers (Supplementary Fig. S7A). Consistent with this observation, the analysis of Oncomine database ^42^ also showed that SPIN1 mRNA expression is extensively upregulated in melanoma tissues when compared with normal skin tissues (∼2.367 folds upregulation, Supplementary Fig. S7B). Moreover, using databases ^40,43,44^ that contain gene expression profiles of clinical cancer samples combined with patient outcomes, we found that overexpression of SPIN1 is correlated with poorer prognosis in patients with breast cancer, colorectal cancer and gastric cancer (Supplementary Fig. S7C-S7F). These data further support that SPIN1 may play an oncogenic role in human cancer progression.

## Discussion

The tumor suppressor p53 provides a critical brake on cancer development in response to ribosomal stress, as impairing this ribosomal stress-uL18/uL5-p53 pathway could accelerate tumorigenesis in c-Myc transgenic lymphoma mice ^45^. However, it remains largely elusive whether this pathway is subjected to the regulation by other yet unknown proteins. In our attempt to understand molecular insights into this possible regulation, we identified SPIN1, the nucleolar protein important for rRNA synthesis ^34^, as a novel regulator of the uL18-MDM2-p53 pathway through interplay with uL18 (Fig. 6E). Our studies as presented here provide the first line of evidence for that SPIN1 acts as an upstream regulator of uL18’s accessibility to MDM2 for p53 activation.

Using IP-MS analysis, we identified SPIN1 as a new uL18-associated protein (Fig. 1A). Our biochemical and cellular experiments using co-IP and GST pull down assays further validated the direct association of SPIN1 with uL18 (Fig. 1B-1D; Fig. 6A-6D). Moreover, we found that SPIN1 and uL18 co-localized in the nucleolus by immunofluorescence assay (Fig. 4D and S3B). Remarkably, SPIN1 specifically binds to uL18, but not uL5 and uL14, as no binding was detected between ectopic SPIN1 and uL5 or uL14 by co-IP (Fig. 1E). Interestingly, SPIN1 does not appear to bind to MDM2, as it was not co-immunoprecipitated with MDM2 either (Fig. 4B). Although our previous reports described a complex of uL18/uL5/uL14-MDM2 ^12^, our present findings indicate that SPIN1 may work in a separate complex with uL18 differed from reported RPs-MDM2 complexes. Also our results suggest that SPIN1 may retain uL18 in the nucleolus so that the latter is unable to shuttle to the nucleoplasm and to inhibit MDM2 activity towards p53 (Fig. 6E).

SPIN1 expression in cells is tightly controlled, as several studies have shown that SPIN1 expression could be negatively monitored by some tumor suppressive non-coding RNAs, such as miR-489 and miR-219-5p ^36,46,47^. Moreover, elevated expression of SPIN1 was strongly correlated with advanced histological stage, chemoresistance and metastasis in patients with breast cancer ^29^. Consistent with the aforementioned oncogenic role of SPIN1, our study as presented here showed that SPIN1 depletion by its specific shRNA leads to the augment of the p53-dependent cancer cell growth arrest and apoptosis. This is at least partly because SPIN1 can promote MDM2-dependent ubiquitination and degradation of p53 (Fig. 3), which is highly likely attributable to its capability to prevent uL18 from binding to MDM2 through retaining uL18 in the nucleolus (Fig. 4E).

Also, knockdown of SPIN1 led to the increase of ribosome-free uL18 and uL5 levels, of the uL18/uL5-MDM2 complex, and of p53 level and activity (Fig. 5A and 5B). The activation of p53 by knocking down SPIN1 is due to the ribosomal stress caused by the depletion of this nucleolar protein, as SPIN1 is required for rRNA synthesis by RNA polymerase I ^34^. Also, consistent with these observations, overexpression of SPIN1 reduced the activation of p53 by Actinomycin D treatment (Fig. 5C), whereas knockdown of uL18 or of uL5 impaired the activation of p53 by SPIN1 knockdown (Fig. 5D and 5E). Several genes have been implicated to modulate the RPs-MDM2-p53 pathway through interplay with ribosomal proteins ^24,25,48-50^. In particular, SRSF1 was identified as a component of the RP-MDM2-p53 complex, and could stabilize p53 via uL18 ^26^. Different from their studies, SPIN1 specifically forms an independent complex with uL18, but not MDM2 or other ribosomal proteins, such as uL5 or uL14, and acts as a negative regulator of p53. Therefore, our present findings unveil a novel mechanism for suppression of the uL18-MDM2-p53 pathway by SPIN1, whose depletion consequently leads to p53-dependent cell growth inhibition and apoptosis.

Consistent with its oncogenic activity, SPIN1 is often amplified in a panel of cancer types with less or no p53 mutation based on our analysis of human samples available in TCGA database (Supplementary Fig. S7A-S7B). In addition, elevated SPIN1 expression correlates with poor prognosis in breast, colorectal and gastric cancer patients (Supplementary Fig. S7C-S7F), further indicating that SPIN1 acts as a potential oncogene. In line with these database observations, we found that overexpression of SPIN1 promotes cancer cell survival, while knockdown of SPIN1 leads to cancer cell death as well as the suppression of cancer cell growth and colony formation predominantly in wild type p53-containing cancer cells (Fig. 2). Remarkably, knockdown of SPIN1 inhibited xenograft tumorigenesis derived from human colon cancer cells, which was much more significantly in HCT116^p53+/+^ cells than in HCT116^p53-/-^ cells (Fig. 7). These results demonstrate that SPIN1 can promote tumor growth and survival by inactivating p53 and its pathway (Fig. 6E).

Intriguingly, we also found that SPIN1 ablation had a moderate inhibitory effect on cell growth in p53-null HCT116 cells in vitro and in vivo as mentioned above (Fig. 2 and 7). These findings suggest that SPIN1 might also possess p53-independent oncogenic effects, which might be explained by two possible mechanisms. First, SPIN1 has been reported to execute its oncogenic potentials by activating Wnt and PI3K/Akt pathways ^29,30^, both of which are closely correlated with cancer progression ^51,52^. Second, since our previous study has demonstrated that uL18 and uL5 could inactivate TAp73 through association with MDM2 ^53^, it is possible that the SPIN1-uL18 interaction might impose suppression on TAp73 activity as well, ultimately leading to cell growth arrest and apoptosis.

Recent studies have demonstrated the role of SPIN1 in rRNA transcription ^32,33^, which provides a clue that dysregulation of SPIN1 may perturb ribosome biogenesis. In fact, in our current study, we observed that SPIN1 deletion per se increases the levels of ribosome-free uL18 and uL5, accompanying elevated p53 protein levels, which recapitulates the effects of ribosomal stress. Our observation that p53 induction caused by SPIN1 depletion could be abrogated by knockdown of either uL18 or uL5 further supports this hypothesis. Therefore, while it is conceivable that SPIN1 counteracts p53 by blocking the interaction between uL18 and MDM2 as discussed above, the mechanism by which disruption of SPIN1 causes ribosomal stress may be also responsible for p53 activation.

In summary, our findings unveil SPIN1 as another novel and important regulator of the MDM2-p53 pathway by predominantly inhibiting the association of uL18 with MDM2 to modulate p53 activity (Fig. 6E) and provide molecular insights into the fine regulation of this pathway as well as a potential target for the future development of an anti-cancer therapy.

## Materials and Methods

### Cell culture and transient transfection

U2OS, H1299, HEK293 and H460 cells were purchased from American Type Culture Collection (ATCC). HCT116^p53+/+^ and HCT116^p53-/-^ cells were generous gifts from Dr. Bert Vogelstein at the John Hopkins Medical institutes. MEF^p53-/-; Mdm2-/-^ cells were generous gifts from Dr. Guillermina Lozano from MD Anderson Cancer Center, the University of Texas. STR profiling was performed to ensure cell identity. No mycoplasma contamination was found. All cells were cultured in Dulbecco’s modified Eagle’s medium (DMEM) supplemented with 10% fetal bovine serum, 50 U/ml penicillin and 0.1 mg/ml streptomycin and were maintained at 37°C in a 5% CO_2_ humidified atmosphere. Cells were seeded on the plate the day before transfection and then transfected with plasmids as indicated in figure legends using TurboFect transfection reagent according to the manufacturer’s protocol (Thermo Scientific, R0532). Cells were harvested at 30-48 h post-transfection for future experiments.

### Plasmids and antibodies

The Myc-tagged SPIN1 plasmid was generated by inserting the full-length cDNA amplified by PCR into the pcDNA3.1/Myc-His vector at EcoR I and Bam HI, using the following primers, forward-CGGAATTCatgaagaccccattcggaaag; reverse-CGGGATCCggatgttttcaccaaaatcgtag. Flag-SPIN1 was generated by inserting SPIN1 cDNA into 2Flag-pcDNA3 at BamHI and XhoI sites. The primers used for PCR amplifying reverse transcribed mRNA were: forward-CGGGATCCaagaccccattcggaaagaca; reverse-CCGCTCGAGctaggatgttttcaccaaatcgta. The GST-tagged SPIN1 fragments, His-tagged SPIN1 and GFP-tagged SPIN1 plasmids were generous gifts from Drs. Bing Zhu from Institute of Biophysics, Chinese Academy of Sciences, and Haitao Li from Tsinghua University, Beijing, China. The plasmids SPIN1 shRNA −1 and −2 were purchased from Sigma-Aldrich (St Louis, MO, USA). The plasmids encoding HA-MDM2, Flag-uL18, Flag-uL5, Flag-uL14, GFP-uL18, p53, His-Ub, GST-MDM2, His-uL18 and GST-uL18 fragments were described previously ^12,19^. Anti-Flag (Sigma-Aldrich, catalogue no. F1804, diluted 1:3000), anti-Myc (9E10, Santa Cruz Technology, catalogue no. sc-40, diluted 1:1000), anti-GFP (B-2, Santa Cruz Technology, catalogue no.sc-9996, diluted 1:1000), anti-SPIN1 (Proteintech, catalogue no. 12105-1-AP), anti-p53 (DO-1, Santa Cruz Technology, catalogue no. sc-126, diluted 1:1000), anti-p21 (CP74, Neomarkers, Fremont, catalogue no. MS-891-P0, diluted 1:1000), anti-PUMA (Proteintech, catalogue no. 55120-1-AP), anti-β-actin (C4, Santa Cruz Technology, catalogue no.sc-47778, diluted 1:5000), anti-GAPDH (Proteintech, catalogue no. 10494-1-AP), anti-tubulin (T6199, Sigma, diluted 1:2000), anti-nucleostemin (H-270, Santa Cruz Technology, diluted 1:1000) were commercially purchased. Antibodies against MDM2 (2A9 and 4B11), uL18 and uL5 were described previously ^12,19^.

### GST fusion protein-protein interaction assay

GST-tagged SPIN1 or GST-tagged uL18 fragments were expressed in *E. coli* and conjugated with glutathione-Sepharose 4B beads (Sigma-Aldrich). His-tagged SPIN1 and His-tagged uL18 were purified using a Ni-NTA (QIAGEN, Valencia, CA, USA) column, and eluted with 0.5 M imidazole. Protein-protein interaction assays were conducted as described previously ^54^. Briefly, for Fig. 6A, 500 ng of purified His-tagged uL18 protein were incubated and gently rotated with the glutathione-Sepharose 4B beads containing 300ng of GST-SPIN1 fragments or GST only at 4°C for 4h. For Fig. 6C, 300 ng of purified His-tagged SPIN1 protein were incubated and gently shaked with the glutathione-Sepharose 4B beads containing 200ng of GST-uL18 fragments or GST only at 4°C for 1h. The mixtures were washed three times with GST lysis buffer (50 mM Tris/HCT pH 8.0, 0.5% NP-40, 1 mM EDTA, 150 mM NaCl, 10% glycerol). Bound proteins were analyzed by IB with the antibodies as indicated in the figure legends.

### Reverse transcription (RT) and quantitative RT-PCR analysis

Total RNA was isolated from cells or tissues using Trizol (Invitrogen, Carlsbad, CA, USA) following the manufacturer’s protocol. Total RNAs of 0.5 or 1.0 μg were used as template for reverse transcription using poly-(T)20 primers and M-MLV reverse transcriptase (Promega, Madision, WI, USA). Quantitative RT-PCR (RT-PCR) was performed using SYBR Green Mix following the manufacturer’s protocol (BioRad, Hercules, CA, USA). The primers for SPIN1, p53, p21, PUMA and GAPDH cDNA are as follows: SPIN1, 5’-CAGAGCTGATGCAGGCCAT-3’ and 5’-ACTGGGTAACAGGGCCATTG-3’, p53, 5’-CCCAAGCAATGGATGATTTGA-3’ and 5’-GGCATTCTGGGAGCTTCATCT-3’; p21, 5’-CTGGACTGTTTTCTCTCGGCTC-3’ and 5’-TGTATATTCAGCATTGTGGGAGGA-3’; PUMA, 5’-ACAGTACGAGCGGCGGAGACAA-3’ and 5’-GGCGGGTGCAGGCACCTAATT-3’; pre-rRNA, 5’-GCTCTACCTTACCTACCTGG-3’ and 5’-TGAGCCATTCGCAGTTTCAC-3’; 18S rRNA, 5’-GCTTAATTTGACTCAACACGGGC-3’ and 5’-AGCTATCAATCTGTCAATCCTGTC-3’; rRNA, 5’-TGAGAAGACGGTCGAACTTG-3’ and 5’-TCCGGGCTCCGTTAATGATC-3’; GAPDH, 5’-GATTCCACCCATGGCAAATTC-3’ and 5’-AGCATCGCCCCACTTGATT-3’.

### Flow cytometry analysis

Cell transfected with scramble shRNA or SPIN1 shRNAs as indicated in the figure legends were fixed with 70% ethanol overnight and stained in 500 μl of propidium iodide (PI, Sigma-Aldrich) stain buffer (50 μg/ml PI, 200 μg/ml RNase A, 0.1% Triton X-100 in phosphate-buffered saline) at 37°C for 30 min. The cells were then analyzed for DNA content using a BD Biosciences FACScan flow cytometer (BD Biosciences, San Jose, CA, USA). Data were analyzed using the CellQuest (BD Biosciences) and Modfit (Verity, Topsham, ME, USA) software programs.

### Cell viability assay

To assess the long-term cell survival, the Cell Counting Kit-8 (CCK-8) (Dojindo Molecular Technologies, Rockville, MD, USA) was used according to the manufacturer’s instructions. Cell suspensions were seeded at 2000 cells per well in 96-well culture plates at 12 h post-transfection. Cell viability was determined by adding WST-8 at a final concentration of 10% to each well, and the absorbance of these samples was measured at 450 nm using a Microplate Reader (Molecular Device, SpecrtraMax M5e, Sunnyvale, CA, USA) every 24 h for 5 days.

### Colony formation assay

Cells were trypsinized and seeded at equal number of cells on 60-mm plates. Media were changed every 4 days until the colonies were visible. Puromycin was added into the media for selection at a concentration of 2 μg/ml. Cells were fixed with methanol and stained with crystal violet solution at RT for 30 min. ImageJ was used for quantification of the colonies.

### Western blot analysis

Cells were harvested and lysed in lysis buffer consisting of 50 mM Tris/HCl (pH 7.5), 0.5% Nonidet P-40 (NP-40), 1 mM EDTA, 150 mM NaCl, 1mM dithiothreitol (DTT), 0.2 mM phenylmethylsulfonyl fluoride (PMSF), 10 μM pepstatin A and 1 mM leupeptin. Equal amounts of clear cell lysates (20-80 μg) were used for WB analysis as described previously ^55,56^.

### In vivo ubiquitination assay

HCT116^p53-/-^ cells were transfected with plasmids encoding p53, HA-MDM2, His-Ub or Myc-SPIN1 as indicated in the figure legends. At 48 h after transfection, cells were harvested and split into two aliquots, one for WB analysis and the other for ubiquitination assay. Briefly, cell pellets were lysed in buffer I (6 M guanidinium-HCT, 0.1 M Na_2_HPO_4_/NaH_2_PO_4_, 10 mM Tris-HCl (pH 8.0), 10 mM β-mercaptoethanol) and incubated with Ni-NTA beads (QIAGEN) at room temperature for 4 h. Beads were washed once with buffer I, buffer II (8 M urea, 0.1 M Na_2_HPO_4_/NaH_2_PO_4_, 10 mM Tris-HCl (pH 8.0), 10 mM β-mercaptoethanol), and buffer III (8 M urea, 0.1 M Na_2_HPO_4_/NaH_2_PO_4_, 10 mM Tris-HCl (pH 6.3), 10 mM β-mercaptoethanol). Proteins were eluted from beads in buffer IV (200 mM imidazole, 0.15 M Tris-HCl (pH 6.7), 30% glycerol, 0.72 M β-mercaptoethanol, and 5% SDS). Eluted proteins were analyzed by WB with indicated antibodies as previously reported ^56^.

### Immunoprecipitation

Immunoprecipitation (IP) was conducted using antibodies as indicated in the figure legends. Briefly, ∼500-1000 μg of proteins were incubated with the indicated antibody at 4 °C for 4 h or overnight. Protein A or G beads (Santa Cruz Biotechnology) were then added, and the mixture was incubated at 4°C for additional 1 to 2 h. Beads were wash at least three times with lysis buffer. Bound proteins were detected by WB analysis with antibodies as indicated in the figure legends.

### RNA interference

SiRNAs against SPIN1, uL18 and uL5 were commercially purchased from Ambion. SiRNAs (20-40 nm) were introduced into cells using TurboFect transfection reagent following the manufacturer’s instruction. Cells were harvested 48-72 h post-transfection for WB or RT-PCR.

### Immunofluorescence staining

Cells were fixed in 4% paraformaldehyde (PFA) for 25 min, followed by permeabilization in 0.3% Triton X-100 for 20 min. The fixed cells were blocked with 5% bovine serum albumin for 30 min, and then the cells were incubated with indicated antibodies at 4°C overnight. Cells were then washed and incubated with the corresponding secondary antibody and 4’-6-diamidino-2-phenylindole (DAPI) for nuclear staining. The cellular localization of SPIN1 or uL18 was examined under a confocal microscope (Nikon, ECLIPSE Ti2).

### Sucrose gradient fractionation and ribosome profiling

This assay was performed following the protocol previously described ^57^. Briefly, cells were harvested at 70-80% confluence after halting translation by 100 μg/ml cycloheximide incubation for 10 min. Cells were lysed in lysis buffer (10 mM Tris-HCl (pH 7.4), 5 mM MgCl_2_, 100 mM KCl, 1% Triton X-100) and gently sheared with a 26-gauge needle for 4 times. Lysates were subjected to 10-50% sucrose gradient centrifugation and the fractions were collected through BR-188 Density Gradient Fractionation System (Brandel, Gaithersburg, MD, USA).

### Generating stable cell lines

Briefly, scramble shRNA or SPIN1 shRNAs purchased from Sigma were transfected into HCT116^p53+/+^ and HCT116^p53-/-^ cells using TurboFect reagent. The cells were maintained at 37°C in a 5% CO_2_ humidified atmosphere for 48 h and were split to two aliquots, one for WB analysis and the other for selection using final concentration of 2 μg/ml puromycin in growth medium.

### Mouse xenograft experiments

Seven-week-old female NOD/SCID mice were purchased from Jackson Laboratories. Mice were randomized into two groups (6 mice in each) and subcutaneously inoculated with 5×10^6^ HCT116 cells that stably expressing scramble shRNA or SPIN1 shRNA in the right and left flanks, respectively. Tumor growth was monitored every other day with electronic digital calipers (Thermo Scientific) in two dimensions. Tumor volume was calculated with the formula: tumor volume (mm^3^) = (length×width^2^)/2. Mice were sacrificed by euthanasia, and tumors were harvested and weighed. To detect p53 activation and apoptosis in vivo, the RNAs and proteins were disrupted from tumors via homogenization in Trizol or RIPA buffer, and then subjected to RT-qPCR and WB analysis. The experiment was not blind and was handled according to approved institutional animal care and use committee (IACUC) protocol (#4275R) of Tulane University School of Medicine. The maximum tumor volume per tumor allowed the IACUC committee is 1.5 cm diameter or 300 mm^3^ per tumor.

### Statistical testing

All in vitro experiments were performed in biological triplicate and reproduced at least twice. The Student’s two-tailed t-test was used to determine mean difference among groups. P<0.05 was considered statistically significant, asterisks represent significance in the following way: *, p<0.05; **, p<0.01. The term “n.s” indicates that no significant difference was found. All the data are presented as mean ± SEM.

## AUTHOR CONTRIBUTIONS

Z.-L.F. conducted most of the studies under supervision of S.-X. Z. and H.L.; J.-M.L. initiated the project and performed initial sets of experiments to conform the uL18-SPIN1 binding and regulation of the uL18-MDM2-p53 pathway, and J.D. performed some plasmids construction, immunofluorescence and IP-WB experiments; B.C. performed part of ribosome-profiling analysis and some plasmids construction and helped in supervising Z.-L.F and editing the manuscript; P.L. and T.L. provided critical reagents and technical assistance, J.-P.X. mentored Z.-L.F.; S.X.Z, and H.L. conceived and designed the study; Z.-L.F. and H.L composed the manuscript.

## ACKNOWLEDGEMENTS

We thank Drs. Bert Vogelstein and Guillermina Lozano for offering HCT116 cells and MEF cells, respectively, Drs. Bing Zhu and Haitao Li for offering plasmids, the Lu lab members and Dr. Hee-Won Park for active discussion and suggestions. Hua Lu and Shelya X Zeng were supported in part by NIH-NCI grants R01CA095441, R01CA172468, R01CA127724, R21CA190775, and R21 CA201889.

## COMPETING INTERSTS

The authors declare that they have no conflict of interest.

